# Viral communities in *Metania* sp. sponge microbiomes with possible effects on CO_2_ fixation

**DOI:** 10.64898/2026.01.21.700432

**Authors:** Carla Patrícia Pereira Alves, Ritam Das, Otávio Henrique Bezerra Pinto, Georgios Joannis Pappas, Ricardo Henrique Krüger, Janina Rahlff

## Abstract

**Background:** Brazilian sponges of the genus *Metania* (phylum Porifera) are filter-feeding organisms from freshwater ecosystems. Here, we explored viral communities of *Metania* sp., their functional role in the sponge and how they differ from those in surrounding water.

**Results:** We identified 1163 viral operational taxonomic units (vOTUs) from sponge tissue and adjacent water, with 555 vOTUs shared across habitats. Viral diversity was higher in sponges than in water, and community composition differed significantly (PERMANOVA, *p* = 0.037). Sponge-associated vOTUs exhibited broad phylogenetic diversity, including deep-branching and unclassified clades, and several exclusively sponge-associated Caudoviricetes. Virus-host predictions revealed 173 interactions, largely with sponge-associated bacteria, supported by CRISPR spacer matches, variant formation in multiple vOTUs across sponge individuals, and a high prevalence of microbial defence systems, particularly restriction-modification, abortive infection, and CRISPR-Cas pathways. Functionally, viral communities carried diverse auxiliary viral genes, including those involved in amino acid and central carbon metabolism, carbohydrate degradation, fatty acid biosynthesis, stress responses (e.g., metacaspase-1), and sulphur cycling. Nine sponge-associated vOTUs encoded carbonic anhydrase (CA), and phylogenomic as well as structural analyses showed strong conservation of CA active sites between sponge viruses, bacterial symbionts, and the sponge host. Protein-level homology searches revealed broad biogeographic distribution of viral CA homologs across global ocean microbiomes, despite limited nucleotide similarity, highlighting deep functional conservation.

**Conclusions:** These findings reveal a phylogenetically diverse and functionally rich viral community associated with freshwater *Metania* sp., characterized by extensive host interactions, diverse defence mechanisms, and auxiliary metabolic capacities. The structural conservation and widespread distribution of viral carbonic anhydrase genes further suggest ecologically significant roles in carbon transformation within freshwater sponges and potentially across aquatic ecosystems.

## Background

Sponges are ancient metazoans of the phylum Porifera that play an important ecological role in the purification, cycling of organic matter, and energy balance of the aquatic environment due to their sessile and filter-feeding habits. In addition to establishing diverse complex ecological relationships with other organisms, symbiosis with microorganisms has an evolutionary link [1, 2]. The evolutionary history of the group suggests that freshwater sponges originated from marine sponges that dispersed to continental environments and colonized them. Brazilian freshwater sponges of the genus *Metania* sp. (*Metaniidae*) are found in rivers and other freshwater bodies across the country [3, 4]. The genus has 11 species described and exhibits a disjunct circumtropical biogeographic pattern in the Neotropical, Afrotropical, Eastern, and Australian regions [5].

Research on sponges, which are also referred to as holobionts, and their permanent microbiome has gained attention for applications in biotechnology (reviewed by de Oliveira, et al. [6]) given the production of biomedically relevant natural products and chemical defences by the sponges [7, 8]. Symbiotic relationships of sponges with a diverse array of microorganisms, including bacteria, archaea, fungi, unicellular eukaryotes, and viruses are common [9, 10]. Sponge symbionts support the sponge metabolism with nutrient cycling and uptake of elements into the sponge tissue [11–13]. In addition, symbionts can act in pathogen defence, and overall host health [14]. Metagenomic studies have revealed different microbial defence systems including restriction modification (RM) systems and adaptive Clustered Regularly Interspaced Short Palindromic Repeats (CRISPR) immunity in marine and freshwater sponge metagenomes [15, 16]. Such defensive systems are commonly used to fend off mobile genetic elements including viruses [17]. They are needed to keep the sponge microbiome in balance by preventing disruptions caused by invading mobile genetic elements (MGEs) and maintaining the stability of symbiotic microbial communities.

While prokaryotic symbionts have been well-studied, the viral component of sponge microbiomes has gained far less attention [18]. With the onset of the era of –omics approaches, studies have revealed that the sponges and seawater both contain a diverse array of viruses (reviewed by Jamal, et al. [19]). These include novel RNA viruses including viruses assumed to infect the sponge [20–22]. In accordance with the fact that most sponge symbionts are heterotrophic bacteria [23], presence of bacteriophages in the microbiome of sponges has also often been reported [24–27]. In addition, nucleocytoplasmic large DNA viruses like Mimiviridae and Poxviridae [26, 28], which can exhibit host-specific and site-specific patterns [26] have been detected. Diseased endemic Lake Baikal sponge *Lubomirskia baikalensis* contained a different DNA viral community compared to healthy individuals. Here, sponge-associated viruses belonged to 16 families being able to infect various organisms [29]. In another Lake Baikal sponge study, related viral scaffolds and virotypes for freshwater sponge *Baikalospongia bacillifera* and marine sponge *Ianthella basta* were detected [15], suggesting that horizontal gene transfer or the presence of closely related viral populations may occur across distinct aquatic environments.

While in marine sponges, the prokaryotic community is not clearly sponge-specific [30], data on the variation of the sponge viral community composition remain limited. Sponge viromes have been shown to respond dynamically to environmental stressors, such as temperature increases [31]. Moreover, lysogeny, i.e., the integration of viral genomes into host genomes, appears to be the dominant viral replication mode within sponge microbiomes, contrasting with the predominance of lytic viruses in surrounding seawater [32]. Sponges were found to have a function in removing marine viruses from the water column and in coral reefs by their filtering activities [33, 34], potentially facilitating the acquisition of (pro)viruses from their aquatic environment [32]. However, it remains unclear whether viruses can be vertically transmitted through sponge developmental stages [14].

Functional studies have begun to shed light on virus-host interactions within sponges. Jahn, et al. [24] identified a phage-encoded ankyrin protein implicated in the sponge’s immune response against bacterial colonization. Although sponges lack a nervous system, they exhibit nervous system-like mechanisms to perceive and react to external stimuli [35]. Sponge-associated bacteria contribute to such neuro-sensory functions; for example, symbionts of the demosponge *Amphimedon queenslandica* produce dopamine and trace amines that act as neurotransmitters and neuromodulators [36]. Additionally, bioactive compounds extracted from sponge-associated microbes have demonstrated antiviral properties [37]. Despite these advances, the role of sponge-associated viruses in mediating host-symbiont communication and environmental adaptation remains largely unexplored. Recent studies have revealed that viruses can encode genes that modulate host or symbiont metabolism, influencing key biogeochemical processes [38, 39]. Among these, viral carbonic anhydrases (CAs), enzymes that catalyze the reversible conversion of CO_2_ and bicarbonate have been identified in marine viruses [40] and microbial biofilms [41]. Investigating whether freshwater sponge-associated viruses encode CA genes may uncover previously unrecognized mechanisms through which viruses shape host-microbe interactions and ecological adaptation.

This study aims to identify and characterize microbial viruses associated with a recently discovered freshwater sponge *Metania* sp. (Figure 1b) through viral metagenomics, and to elucidate the functional roles of virus-encoded genes in host-microbe interactions. Understanding the diversity, distribution, and ecological functions of the *Metania* sponge virome benefits our understanding of virus-prokaryote-sponge interactions and supports the conservation of higher organisms inhabiting these environments in Brazil, particularly under increasing pressures from human-driven habitat destruction, pollution, and global (climate) change.

**Figure 1:**
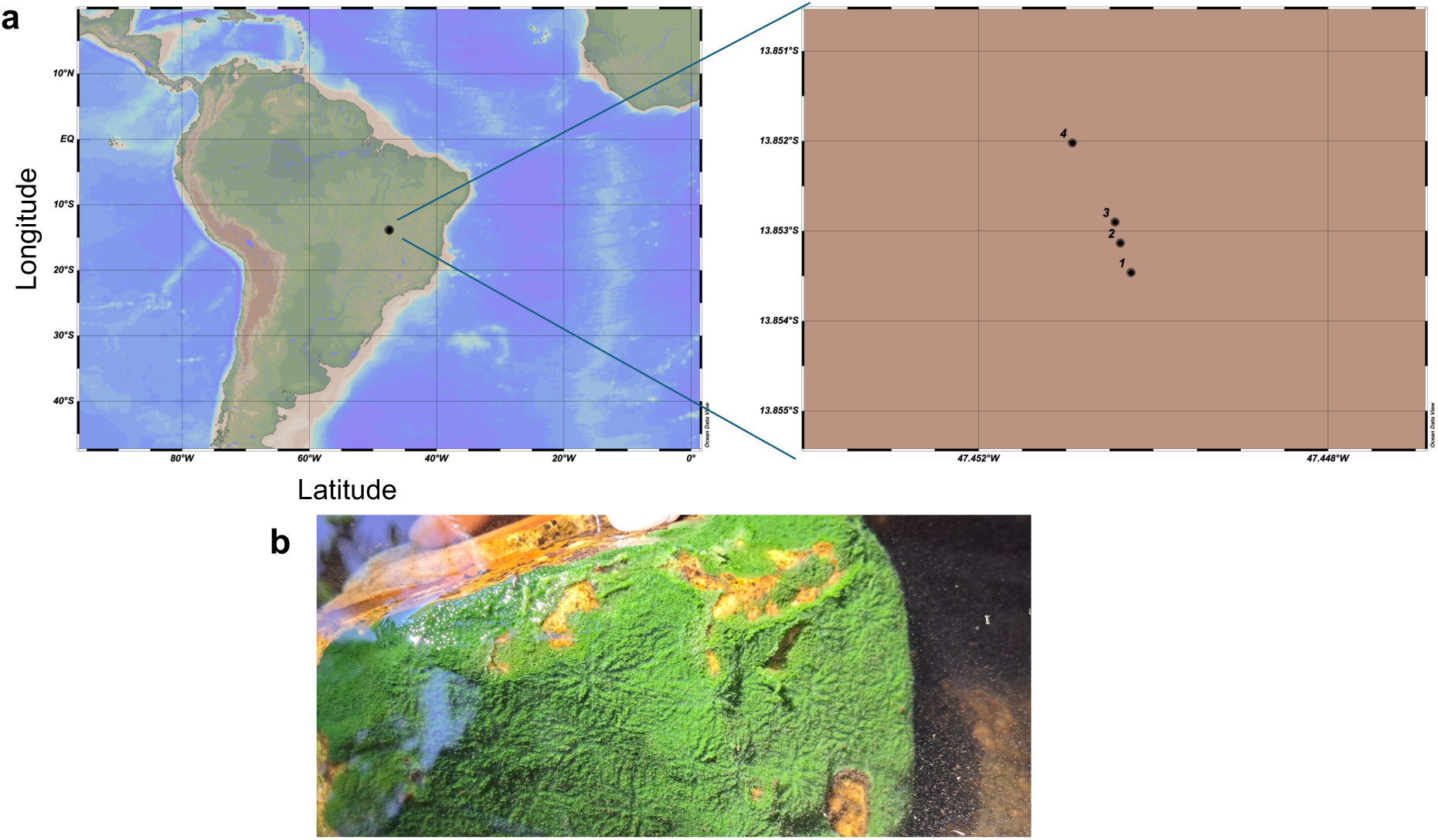
Sampling location and sponge sample. a) Map of the four sampling sites in Brazil, created with Ocean Data View v.5.8.2. [108]. Depicted sampling stations 1 – 4 match water and sponge samples 1 – 4. b) Image of the sponge *Metania* sp. attached to a rock (Photographer: Rafaella Silveira).

## Results

### Viral diversity in sponge and surrounding water

We identified 1163 viral operational taxonomic units (vOTUs) >10 kb genome size, of which 649 and 262 were predicted to belong to Caudoviricites as predicted by GeNomad and vConTACT2, respectively (Table S1 & 2). From the 649 vOTUs, 548 and 320 vOTUs assigned to Caudoviricetes were detected in water and sponge samples, respectively, and 216 shared between both habitats. In the sponge samples, the number of unclustered vOTUs (*M* +/− STD = 65.5 % ± 7.7, Figure 2a) was higher than for water-derived vOTUs (*M* = 39.9 % ± 0.95). The viral cluster VC_833 was detected in all samples and showed the highest relative abundance in water (mean = 20.2 % ± 0.6). Protein-based clustering identified its closest neighbour as Thermoanaerobacterium phage THSA-485A. Next most abundant was by VC_864_0 (*M* = 9.9 % ± 0.6), which has proteins related to Microcystis phage Mae-Yong1326-1. The Shannon-Wiener diversity index ranged from 5.6 – 5.8 and 5.1 – 5.3 for sponge and water viral communities, respectively (Figure 2b). Non-metric multidimensional scaling (NMDS) analysis revealed distinct clustering of sponge and water viral communities (Figure 2c). A PERMANOVA confirmed that the sponge-water distinction significantly influenced community composition (R² = 0.96, F = 145.82, *p* = 0.037), explaining 96 % of the observed variance. Looking at the dispersion of viral communities within each group based on their distance to the group centroid, sponge samples show greater variability in community composition, indicated by a wider spread of distances, whereas water samples have less dispersion, reflected by a more compact distribution (Figure 2d). Overlaps between vOTUs in water and sponge were detected as well: in total 555 vOTUs were found in both water and sponge tissue, 140 vOTUs were only sponge associated, and 468 vOTUs were specific to the surrounding water (Figure 2e).

**Figure 2:**
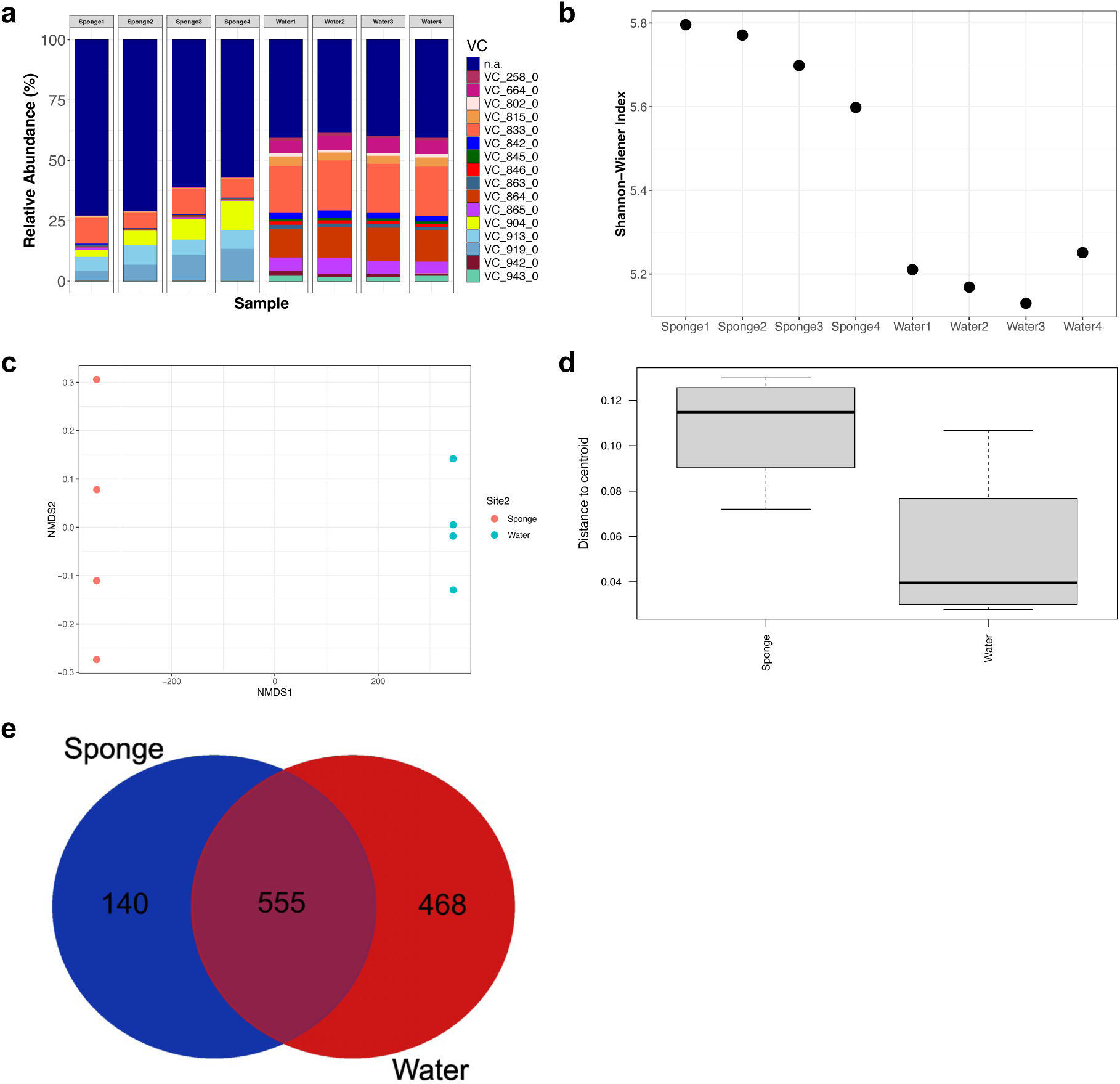
Viral diversity analysis. Relative abundance of top50 vOTUs forming viral clusters (VC) (a) Shannon-Wiener diversity between sponge vs water samples (b) NMDS plots showing community distribution (c) corresponding box plots showing multivariate dispersion for the variation in viral community composition within each group (d). Venn diagram showing overlapping vOTUs (e).

### Proteomic tree reveals broad phylogenetic placement of vOTUs

To explore the taxonomic diversity of sponge– and seawater-associated viruses, we built a ViPTree proteomic tree using normalized tBLASTx scores, incorporating 120 vOTUs from our metagenomes and reference viral genomes (Figure 3). Out of the 120 vOTUs determined as complete, high, and medium-quality based on CheckV (Table S3), two sponge-related viruses formed a protein-based clade with *Bracoviriform inaniti* (comb-NC_005165), an insect virus. Additionally, 44 of the high quality vOTUs, found in either water and sponge or sponge only, clustered closely with this virus, suggesting they may also represent eukaryotic viruses, potentially sponge viruses. Eight vOTUs predicted as Caudoviricites were only found on the sponge tissue but not in the surrounding water. Sponge-associated vOTUs were distributed across a wide range of clades, many of which fall within the class Caudoviricetes, which encompasses tailed bacteriophages. Several sponge-derived vOTUs clustered within well-characterized viral lineages, while others occupied deep-branching or unclassified clades, indicating substantial genomic divergence from existing references. Some vOTUs formed monophyletic groups lacking close reference genomes, which may represent previously undescribed viral genera or families. Meanwhile, others grouped with reference genomes isolated from diverse aquatic environments such as Winogradskyella phage Peternella_1 (NC_062754) from the North Sea [42], Pantoea phage PdC23 (NC_071008) from a Tunisian wastewater system [43], or Bacteriophage DSS3_VP1 (NC_055826) from waters surrounding Venice, Italy [44]. The reference phage Vibrio phage CHOED (NC_023863), isolated from molluscs [45], clustered with vOTUs from this study, suggesting that viruses typically associated with animal hosts may share evolutionary origins or ecological niches with free-living viral populations, indicating potential connectivity between host-associated and environmental viral communities. The phylogenetic breadth of sponge-derived vOTUs overall highlights the ecological complexity and evolutionary novelty of viral communities associated with sponges.

**Figure 3:**
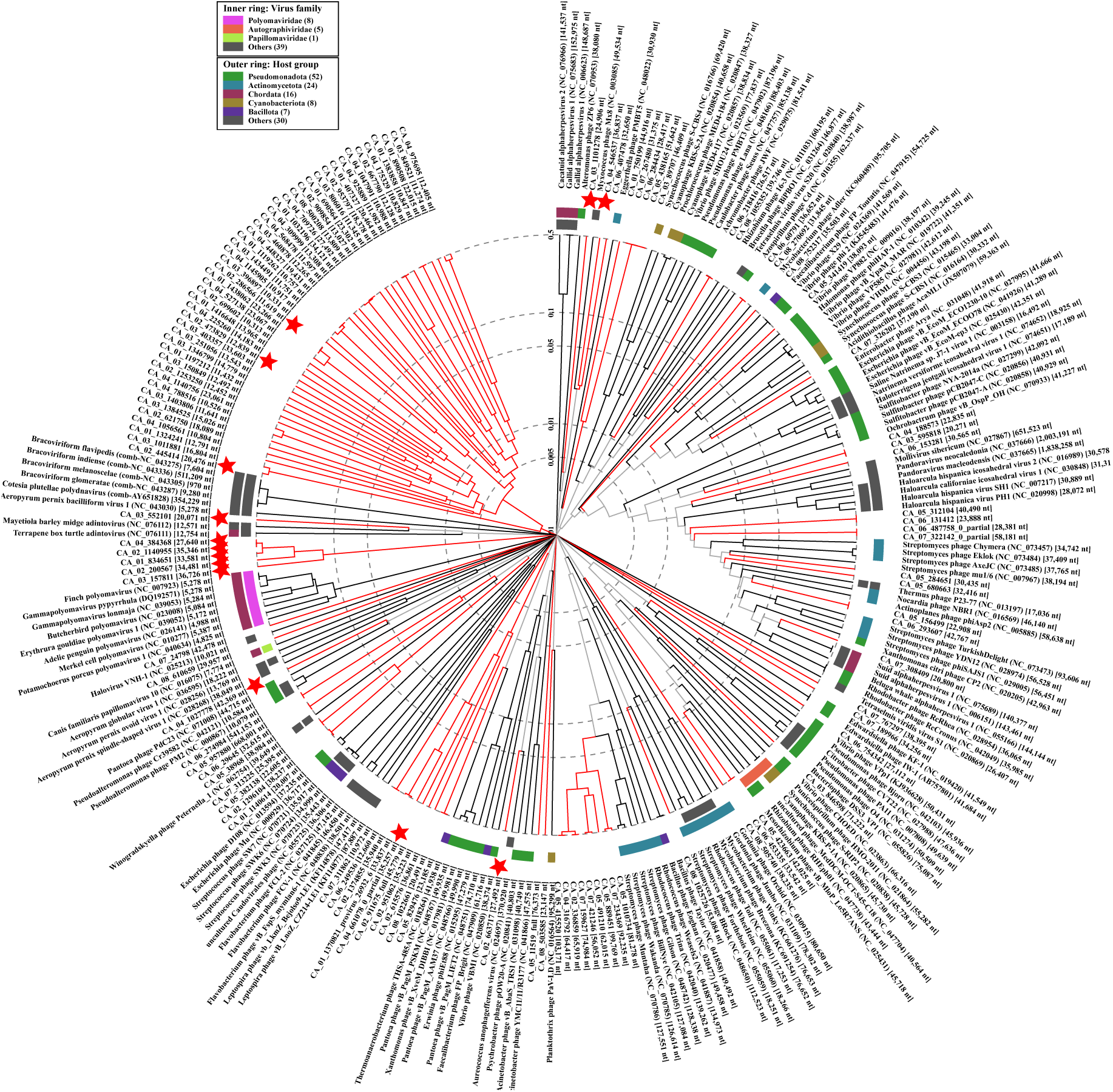
Viral proteomic tree for 120 vOTUs clustering with reference viruses. Red stars denote vOTUs detected in sponges only (based on read mapping), while red lines highlight clades that contain at least one vOTU (from sponge or seawater).

### Virus-host associations in the sponge microbiome

In total, 173 vOTU-host interactions were predicted for 148 unique vOTUs (Table S4). The vOTUs were assigned to different *Burkholderiaceae* genera (*Comamonas, Jezberella, Hylemonella, Limnohabitans_A, Hydrogenophaga, Pelomonas, Polynucleobacter, Rhodoferax_C, Xylophilus*), members of the family *Sphingomonadaceae* (*Novosphingobium, Sandarakinorhabdus),* the family *Chitinophagaceae (Sediminibacterium, Hanamia),* family *Nostocaceae (Dolichospermum),* family *Methylomonadaceae (Methylobacter_A, Methyloglobulus),* members of the families *Xanthomonadaceae* and *Pseudomonadaceae (Pseudomonas, Stutzerimonas, Stenotrophomonas, Xanthomonas)* among others (Figure 4a). The composition and function of the microbial community will be addressed elsewhere [46].

**Figure 4:**
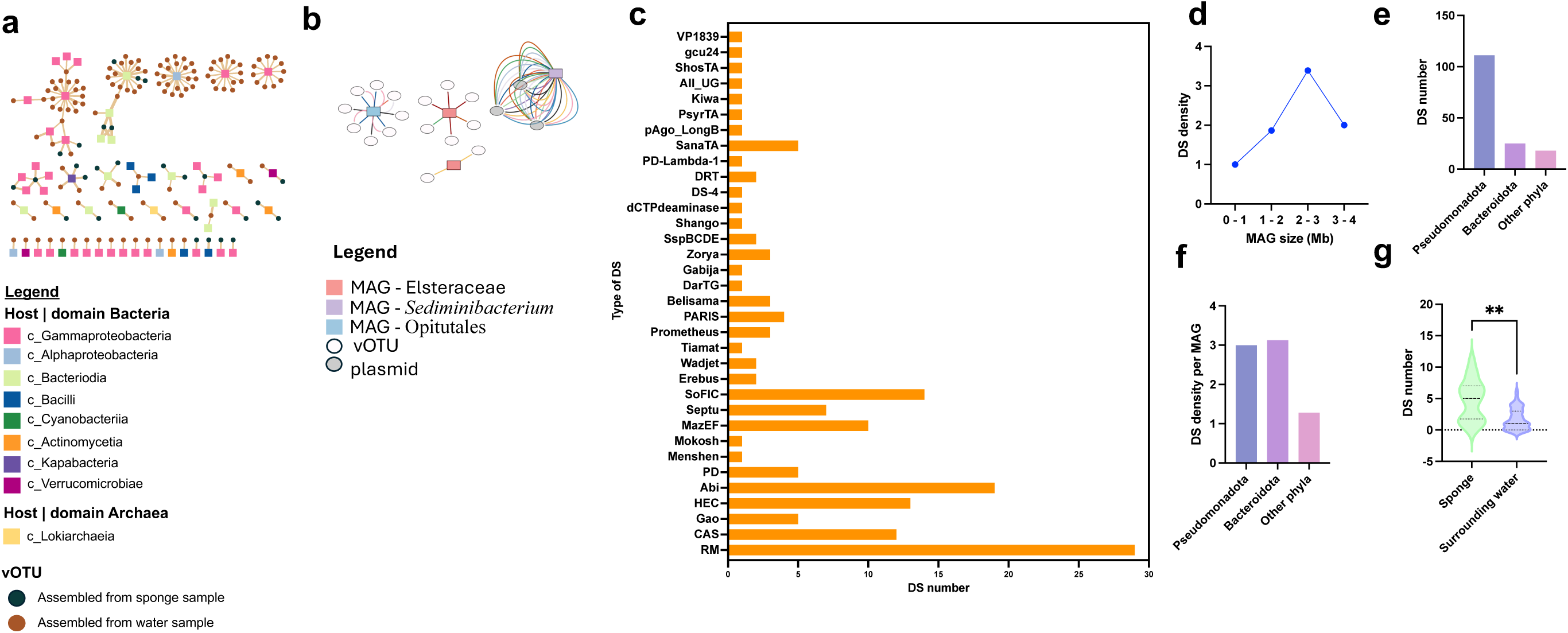
Host assignment to vOTUs and CRISPR-based interactions. Host prediction network for all viral operational taxonomic units (vOTUs) that had a predicted prokaryotic host (a). CRISPR-targeting of vOTUs and plasmids (b). Here, same spacer (=line) color indicates the same spacer sequence if derived from the same metagenome assembled genome. Prevalence of diverse defense systems in the sponge and surrounding water MAGs. (c). DS density was found to be enriched in genomes of 2 – 3 Mb size (e). Comparison of DS number among Pseudomonadota, Bacteroidota, and other phyla (f). Comparison of DS density (per MAG) among Pseudomonadota, Bacteroidota, and other phyla (g). Comparison of DS number among MAGs associated with sponges and surrounding water indicated sponge MAGs to encode significantly higher DS.

Three metagenome-assembled genomes (MAGs) contained CRISPR spacers matching vOTUs. Notably, a sponge-associated MAG assigned to an *Elsteraceae* bacterium (96.5% completeness, 2.2% contamination, Table S5) had four distinct CRISPR spacers from two CRISPR arrays that matched eight sponge-derived vOTUs found across different sponge samples (Table S6). Read mapping showed that none of these eight vOTUs were present in the surrounding water, suggesting that the CRISPR targeting of these phages is specific to the sponge environment. A freshwater-derived MAG assigned to an Opitutales bacterium (98% completeness, 0% contamination) contained five CRISPR spacers that matched eight distinct vOTUs originating from both water and sponge sources. In this case, multiple spacers matched the same vOTU (Figure 4b), suggesting potential redundancy in CRISPR targeting or a strong selective pressure to defend against that phage. A water-derived MAG assigned to *Sediminibacterium* (97.9% completeness, 0% contamination) targeted three distinct freshwater-associated plasmid sequences within the sample, each with twelve different CRISPR spacers (Figure 4b) probably indicating that the CRISPR-Cas system may primarily function in defense against plasmid-borne genetic elements rather than phages.

A total of 34 different types of defense systems (DS) were detected among 156 DS in the MAGs. Among the different types, the restriction modification (RM) system was found to be the most common (29) (Figure 4c) followed by the group of abortive infection (Abi) systems (19), the FIC domain-containing system SoFIC (14), and the group Hma-Embedded Candidates (HEC) (13) (Figure 4c). We also found that the DS were prevalent in MAGs with a genome size within the range of 2 – 3 Mb (Figure 4d). A comparison across different phyla indicated that phylum Pseudomonadota (n = 37) had more DS compared to Bacteroidota (n = 7) and other combined phyla (n = 14) (Figure 4e). When normalized by MAG count, DS density was higher in Bacteroidota than in other phyla, especially for the most prevalent DS types. Although RM and CRISPR-Cas systems were more numerous in Pseudomonadota, their density per MAG was higher in Bacteroidota, indicating DS are more consistently represented across its MAGs (Figure 4f, Figure S1), highlighting the potential ecological or evolutionary significance among these microorganisms. Notably, a comparison of DS densities between sponge-associated MAGs and those from surrounding water revealed a significantly higher (*p* < 0.01) and more consistent distribution of DS in sponge-associated MAGs (D’Agostino & Pearson test was performed to analyse normality, followed by a Welch’s t-test, (*p* = 0.0014) (Figure 4g).

### Viruses form variants across sponge samples

We identified three vOTUs with variant formation across the four sponge samples (Table S7). One was a prophage (CA_01_1370821_provirus) related to Caulobacter phage RW, with variants suggesting active evolution and a temperate lifestyle. Another (CA_02_911675), encoding carbonic anhydrase, showed sponge-specific variants, implicating roles in carbon metabolism or pH regulation. The third (CA_04_661078) also formed variants across all samples and was identified as a phage. All three clustered within VipTree, lacked host predictions, encoded phage hallmark proteins, and notably carried Type III secretion system proteins. The first two were additionally detected in water samples, suggesting water-mediated dispersal between sponge hosts.

### Auxiliary viral genes (AVGs) highlight viral strategies to modulate host metabolism

Across the full vOTU set, auxiliary viral genes (AVGs, a new overarching term for auxiliary metabolic genes (AMGs) recently introduced [47]) involved in amino acid metabolism were among the most abundant, including 68 linked to biosynthesis and 38 to degradation (Table S8). We also identified 641 viral tRNA genes that may offset host codon bias and enhance viral protein synthesis. Viral AVGs were enriched in core energy pathways such as C1 metabolism (34), glycolysis (32), the TCA cycle (20), and oxygen-dependent metabolism (9), with additional genes for photosynthesis (4) and methane-based C1 metabolism (2), indicating modulation of host energy systems (Figure 5a). Carbon metabolism AVGs included the pentose phosphate pathway (22), pyruvate metabolism (3), and sugar degradation (galactose, fucose, lactose, xylose), as well as hydrocarbon degradation (4) and the methylaspartate and urea cycles (5 each), reflecting niche-specific adaptations. CAZy-related AVGs included abundant Glycoside Hydrolases (GH23, GH24, GH25, GH73, GH108) for peptidoglycan and chitin degradation, and Glycosyltransferases (GT10, GT11, GT17, GT41, GT60) contributing to glycan modification and adhesion (Figure 5c). Additional enzymes (AA2, AA3, CBM25, CE14, CE4, PL4) supported lignin, starch, xylan, chitin, and pectin degradation, enhancing nutrient acquisition. Class II AVGs targeted indirect host modulation, with peptidases (389) and inhibitors (13) suggesting control of proteolysis. Transport-related AVGs included systems for ions, amino acids, sugars, and specialized substrates, while others linked to motility, adhesion, and cell division imply structural manipulation (Figure 5b). Electron transport (21) and cofactor biosynthesis genes (cobalamin, siroheme, menaquinone) further point to viral support of host energy and redox metabolism.

**Figure 5:**
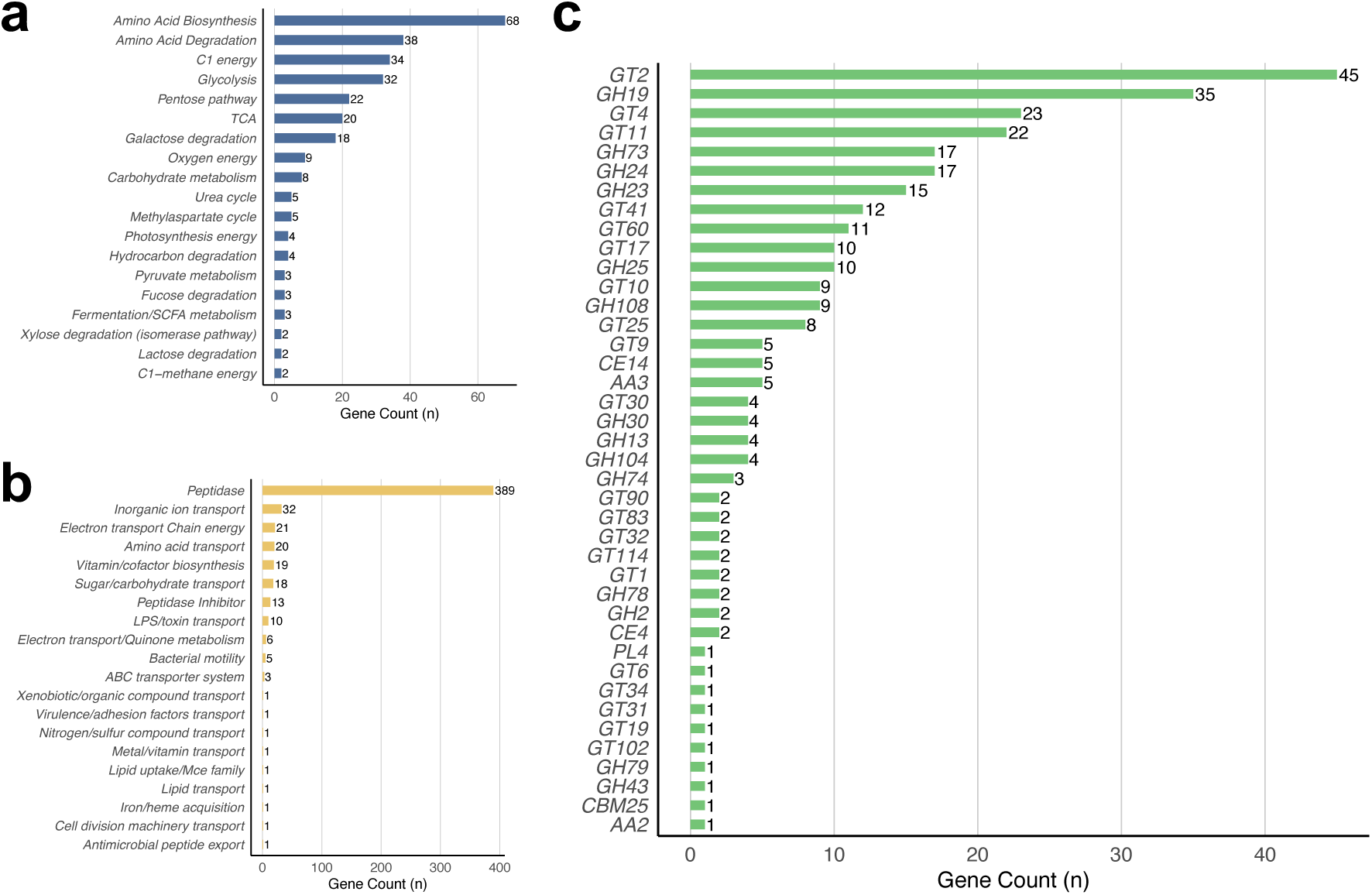
Auxiliary Viral Genes (AVGs). Class I AVGs: Directly involved in core host metabolism, including amino acid biosynthesis and degradation, carbon and energy metabolism, nucleotide synthesis, and other central pathways (a). Class II AVGs: Indirectly involved in the modulation of host metabolism, including transporters, peptidases, motility genes, and the biosynthesis of cofactors and vitamins (b). CAZyme-associated AVGs, grouped by CAZy families (c).

Across the full set of 1163 vOTUs, four vOTUs encoded a β-ketoacyl synthase N-terminal domain (pfam db, PF00109.32) and C-terminal domain (pfam db, PF02801.28). This is an enzyme involved in fatty acid biosynthesis, which has previously been identified in the microbiome of the marine sponge *Arenosclera brasiliensis* [48]. In addition, nine vOTUs carried ORFs functionally annotated as CA (KEGG db, EC:4.2.1.1), suggesting a potential role in host-associated carbon metabolism or pH regulation and further elucidated below. Five vOTUs encoded metacaspase-1 (KEGG database, EC 3.4.22.-), a protease commonly linked to programmed cell death and stress response pathways. CA and metacaspase-1 may best be categorized in the Auxiliary Physiology Genes (APG) category of AVGs [47]. One vOTU (CA_04_1027778) from the sponge tissue but not in water encoded for a phosphoadenosine phosphosulfate (PAPS) reductase, an AVG commonly found in environmental viruses [49] and involved in sulfur metabolism, specifically in the reduction of PAPS to sulfite, which is a crucial step in assimilatory sulfate reduction and the biosynthesis of sulfur-containing compounds.

### Structural and functional conservation of carbonic anhydrase

Functional annotations of MAGs and vOTUs revealed the presence of CA encoding genes. These metalloenzymes in bacteria contribute to the regulation of pH, photosynthesis, and the transportation of carbon dioxide and bicarbonate for other metabolic processes [50]. We identified nine vOTU-encoded CAs, which could essentially function as APGs. To further understand the biological and functional conservation of the proteins, we conducted a phylogenetic analysis of CA sequences from all the bacterial MAGs, vOTUs and the *Metania* sp., followed by structural characterization of these proteins. In the unrooted phylogenetic tree, we observed clear differentiation and formation of three major clades, indicating ancestral relationships that diverged possibly due to sequence differentiation and functional adaptations (Figure 6a). Domain prediction of these sequences indicated that these divergences are a result of distinct clade formation based on the CA family. We observed the MAGs to encode three different genetic families, the α-, β-, and γ-classes of CA (Figure 6a). The β-class of CA was prevalent in the bacterial MAGs and exclusive within the vOTUs and other phages. The CAs from the vOTUs showed a similar ancestry to those of the bacterial MAGs, suggesting a common origin or acquisition through horizontal gene transfer events. This was corroborated by the sequence alignment analysis, which demonstrated the conservation of several key residues in the CAs from the MAGs and the viruses (Figure 6b).

**Figure 6:**
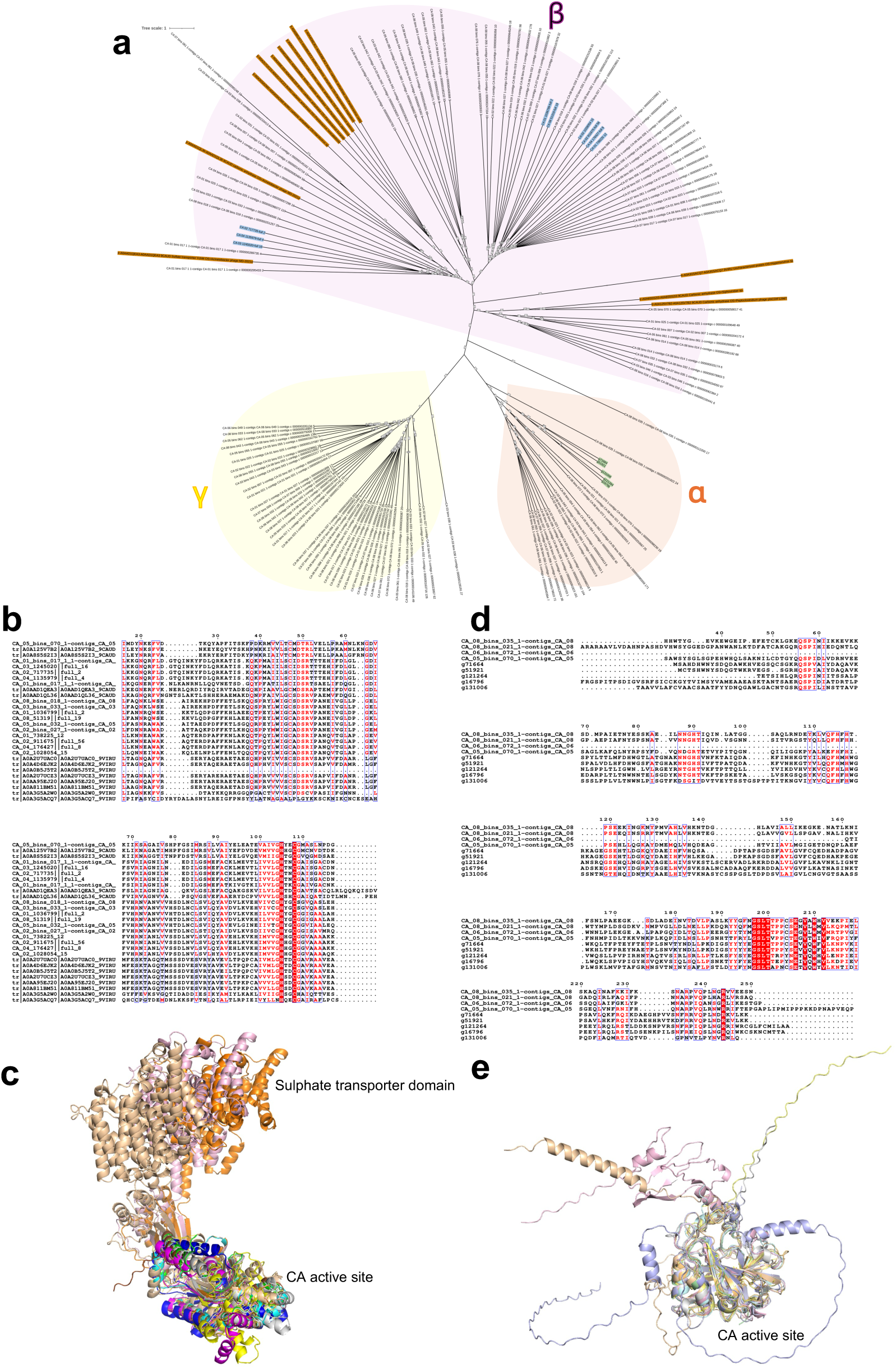
Phylogenetic, sequence, and structural characterization of carbonic anhydrases (CA) derived from the sponge *Metania* sp., bacterial MAGs, and viral OTUs. Phylogenetic analysis of CAs from the *Metania* sp. (green), the bacterial MAGs (no color), vOTUs (blue), and related phages (brown) indicates distinct clade formation based on α- (orange), β- (magenta), and γ-(yellow) classes (a). Multiple sequence alignment of CAs from MAGs, vOTUs and related viruses sharing the same clade in the phylogenetic tree (b). Structural alignment of β-class CAs from MAGs, vOTUs and related viruses sharing the same clade in the phylogenetic tree indicates a conserved CA active site core (c). Accession IDs, amino acid sequences, confidence of structural prediction, RMSD scores and colour codes are listed in Table S9. Multiple sequence alignment of α-class CAs from the bacterial MAGs and *Metania* sp. sharing the same clade in the phylogenetic tree (d). Structural alignment of α-class CAs from MAGs, and *Metania* sp. sharing the same clade in the phylogenetic tree indicates a conserved CA active site core (e). Accession IDs, amino acid sequences, confidence of structural prediction, RMSD scores and colour codes are listed in Table S10. Phylogenetic placement of *Metania* CA genes is given in Figure S3.

The structural and domain analysis revealed multi-domain proteins in some vOTUs, where the CA domain was accompanied by an N-terminal sulphate transporter domain, indicating adaptation of CA to different functional requirements. However, an alignment analysis of high-confidence models (Table S9) indicated structural conservation of the CA enzyme’s active site. Although sequence features of the CAs are noticeably different from each other, the common CA core enzyme-active domain from the viruses (vOTUs and database phages) could be superimposed on the structures from that of the MAGs, with overall low root-mean-square deviation (RMSD) scores ranging between 0.48 – 11.75 (Figure 6c). This low RMSD indicates a high degree of structural conservation of the CA core domain across different sequences, despite their varied functional roles and additional domains. Our analysis highlights the structural and functional diversity of CAs within the sponge vOTUs and the MAGs, emphasizing their conservation despite sequence divergence. This plausibly suggests that CAs may have originated in bacteria and were subsequently transferred to other organisms, such as viruses, through gene transfer events.

In the phylogenetic tree, we also observed distinct clade formation of α-class CAs from bacterial MAGs and from the *Metania* sponge, sharing a common ancestry. Multiple sequence alignment indicated differences between the sponge and MAGs (mainly assigned to phylum Pseudomonadota), but overall conservation of active-site residues (Figure 6d). Structural alignment of high-confidence models (Table S10) of α-clade CA’s from *Metania* and bacterial MAGs also indicated conservation of the active site region in these metalloenzymes (Figure 6e), with overall low RMSD scores ranging between 0.47 – 1.32. Given the shared ancestry, presence of eukaryotic-like CA domains in the bacterial MAGs, conservation of active site residues and structure, the exchange of α-class CAs between *Metania* and bacterial MAGs is highly probable. This eukaryotic-prokaryotic mobility is further supported by the presence of N-terminal intrinsically disordered regions (IDRs) in the CAs encoded by both the bacterial MAGs and *Metania* (Figure 6e). These disordered regions have also been demonstrated in α-class CAs of humans [51], where these regions in the tumour-associated hCA IX comprise a signal peptide and a proteoglycan-like domain potentially acting as an interacting surface.

### Viral carbonic anhydrase genes exhibit conserved protein-level, but not nucleotide-level, homology across diverse marine taxa and geographic locations

To explore the distribution of viral CAs (Table S11) in the global oceans, representative sequences from the three CA clades were queried using BLASTn and BLASTp against the Ocean Gene Atlas (metagenomic dataset OM-RGCv1). At the nucleotide level (BLASTn), CAs from two of the three clades showed no detectable similarity to sequences in OM-RGCv1. Only the CA of vOTU CA_03_1245020||full16 exhibited a single BLASTn hit, with low abundance (57 reads per million total reads, according to OGA normalization) and limited geographic distribution. In contrast, BLASTp results revealed widespread protein-level homology across all CA sequences. Notably, the CA of vOTU CA_03_1245020 produced 4589 hits, with a cumulative normalized abundance of 536755 reads per million total reads (according to OGA normalization), indicating a high level of functional conservation (Figure 7). The remaining sequences also returned between 1238 and 1247 hits, with normalized abundances ranging from 35477 to 37609 reads per million total reads. Protein homologs for the CA of vOTU CA_03_1245020||full16 were taxonomically diverse, spanning Cyanobacteria, Pseudomonadota, and Bacteroidota, highlighting the broad phylogenetic conservation of this CA variant in marine microbial communities (Figure S2).

**Figure 7:**
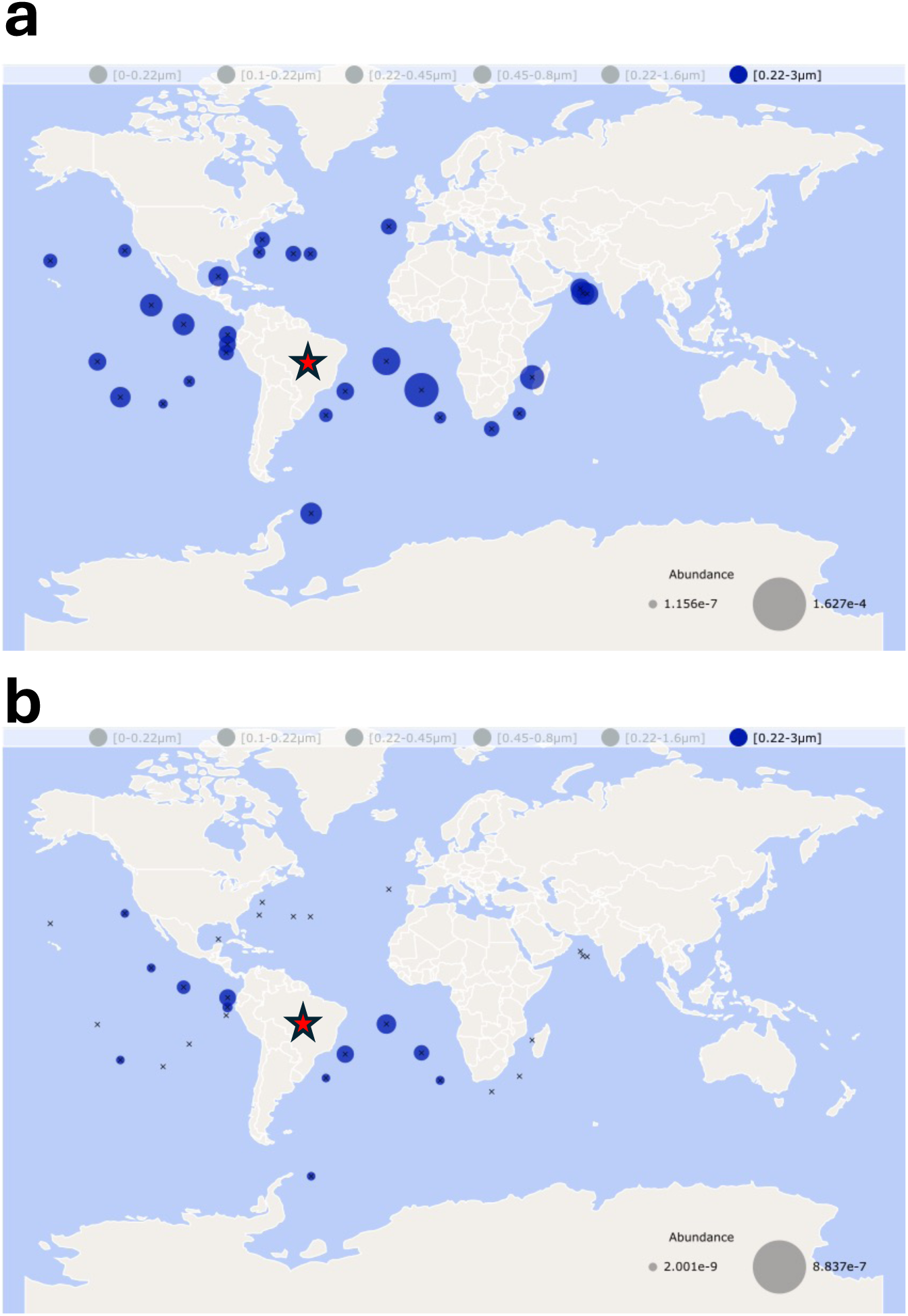
Distribution of vOTU-encoded carbonic anhydrase in the oceans. Geographic distribution of the most abundant viral-encoded CA (CA_03_1245020) based on BLASTp (a) and BLASTn (b) analysis. Abundances were expressed as fractions of total reads per sample. The red star on the world maps indicates the sampling location.

## Discussion

Our study demonstrates that the viral communities associated with *Metania* sp. sponges are significantly distinct from those in the surrounding freshwater environment, both in taxonomic composition and ecological characteristics. This emphasizes that sponges harbour a unique virome likely shaped by host-specific factors. The surrounding water had more viruses classified within the Caudoviricetes, a group of double-stranded DNA bacteriophages that are well-characterized and commonly found in aquatic environments and are most represented in public databases [52]. In contrast, sponge-associated samples contained a markedly higher proportion of unclassified viruses, many of which were entirely absent from water samples. This suggests that these viruses may represent sponge-specific viral lineages, which are potentially adapted to the sponge microenvironment and its dense, host-associated microbial communities. The ecosystem where the sponge originates from can be best described as a shallow, pristine river that runs through an environmental conservation area (Chapada dos Veadeiros National Park) and is used only for ecotourism activities.

We detected CRISPR spacers in a sponge-associated microbial MAG that match vOTUs found exclusively in the sponge, pointing towards an evolutionary history of virus-host interactions taking place within the sponge tissue. CRISPR spacer with viral protospacer interactions have been used to uncover that virus-host interactions occur rather within-communities of different cold-water sympatric sponges compared to between communities [53]. Horn, et al. [16] found that CRISPRs from the microbiome of Mediterranean sponge species had mostly unknown targets but also matched plasmids and viruses. We observed an instance where the same CRISPR spacer matched multiple distinct vOTUs in the sponge microbiome. The reuse of a single spacer may suggest the dynamic co-evolution between microbial hosts and their viruses and thus represents an efficient adaptive immune strategy. Such conservation could also mean that these viruses may belong to closely related lineages or share common evolutionary pressures that preserve specific sequence motifs. From the host’s perspective, the retention and reuse of these spacers may provide broad-spectrum immunity, allowing a single spacer to defend against multiple viral threats. This pattern supports the idea that CRISPR-Cas systems in sponge-associated microbes may be selectively maintained to target ecologically dominant or recurrent viral groups. Our study also identified several types of DSs within the MAGs, with RMS being the most common, as in most microbiomes [54]. Interestingly, sponge-associated MAGs exhibited significantly higher and more consistent DS distributions than those from the surrounding water, where we found more viruses, which suggests an interaction-dependent evolution of the microbiome.

The identification of several AVGs in sponge-associated vOTUs suggests that viruses may play an active and multifaceted role in shaping host-microbe interactions within sponge holobionts and influence host physiology and microbial community structure through targeted metabolic reprogramming and modulation of host stress responses. Viral-encoded metacaspase-1 may participate in controlling host cell fate via programmed cell death [55, 56]. A study by Jiang, et al. [57] analysed metacaspases from cyanobacteria and found two major families (α and β) and various domains involved in signal transduction, with only few playing a role in apoptosis. The presence of CA genes, which have been found in marine sponge-associated viromes before [58], suggests viral involvement in carbon cycling and pH regulation within sponge tissues and freshwater ecosystems, potentially contributing to microenvironmental stability. Members of the α-CA gene family have been shown to be essential for calcium carbonate deposition in calcareous sponges [59]. However, α-CA was also detected in *Metania*, whose siliceous rather than calcitic spicules [60] indicate that the enzyme likely fulfills additional roles within the sponge. We identified three distinct CA gene clades showing strong protein-level homology to marine bacterial sequences but little or no nucleotide similarity. This discrepancy indicates that while viral CA genes are genetically divergent, their encoded proteins remain functionally conserved. Notably, the CA variant vOTU CA_03_1245020||full16 displayed high protein homology to distant marine microbes but shares nucleotide similarity only with a Flavobacteriales strain. Protein homologs of other viral CAs were similarly widespread, primarily among Pseudomonadota and Bacteroidota across multiple oceanic regions. Bacteria are well known to encode a diverse array of CA classes and could theoretically provide this function for their host in symbiotic contexts [61]. Their distribution across diverse hosts supports potential horizontal gene transfer and a role in host carbon metabolism, possibly as APG. Functional conservation despite sequence divergence may reflect adaptation to environmental conditions. The geomorphology of the Chapada dos Veadeiros region, shaped by marine processes about 1.5 billion years ago through the deposition of an ancient epicontinental sea later metamorphosed into quartzite and related lithologies[62], further supports a potential evolutionary link between marine and freshwater systems. Together, these findings suggest that viral CA genes, although originating in freshwater viruses, encode conserved, ecologically relevant functions likely acquired from marine bacteria during evolution. We propose that CA genes may have originated in one habitat and diversified in the other while preserving their catalytic role across ecological boundaries. As key mediators of CO_2_-bicarbonate interconversion, CAs underpin primary productivity and global carbon cycling. In the context of environmental change, viral-encoded CAs could help buffer stress, sustain carbon fluxes, and maintain ecosystem stability under fluctuating pH and CO_2_ conditions. Their broad phylogenetic and environmental distribution also points to recurrent horizontal transfer, potentially from bacteria to viruses (β class) and from bacteria to sponges (α class), where they may have been co-opted for host-specific carbon and mineral metabolism.

## Methods

### Sample collection

DNA samples were collected from four different sites within waterfall complex Veredas, located in Cavalcante, Goiás State, Brazil (Figure 1a). These samples included both specimens of freshwater sponges and water from their surrounding environment. At each location, individuals of the newly identified sponge species *Metania* sp. were gathered by scraping submerged rocks. Water samples (500 mL) were concentrated through 0.22 µm pore size nitrocellulose membranes (Millipore, Burlington, MA, USA) and frozen on dry ice. DNA extraction was conducted using commercial extraction kits: the FASTDNA Spin for Soil kit (MP Biomedicals, São Caetano do Sul, Brazil) for sponge samples and the DNeasy PowerWater Kit (QIAGEN, Hilden, Germany) for water samples, with a protocol modified to incorporate additional thermal cell lysis steps. The extracted DNA was evaluated for quality and concentration using agarose gel electrophoresis and a Qubit fluorometric quantification Q32857 (Invitrogen). Library preparation was done using the TruSeq Nano DNA Kit with TruSeq Nano DNA Sample Preparation Guide, Part # 15041110 Rev. D (Illumina, San Diego, CA, US) by Macrogen (Seoul, South Korea) with subsequent metagenomic sequencing using the Illumina NovaSeq6000 system, producing 150 bp paired end reads.

### Viral metagenomic analysis

Paired-end Illumina sequencing data were initially processed to remove adapters and filter low-quality bases using BBtools v.39.03 (https://sourceforge.net/projects/bbmap/[63]) with a minimum base quality threshold and a minimum read length of 15 (each). High-quality reads were then assembled into contigs using MEGAHIT v 1.2.9 [64], with default parameters. Viral scaffolds larger than 5 kb were identified by using VirSorter2 v.2.2.3 [65] with the setting –-include-groups “dsDNAphage,ssDNA,NCLDV,lavidaviridae”, GeNomad v.1.5.2 with database v.1.3 [66], and VIBRANT v.1.2.1 [67]. Viruses identified from these tools were pooled, filtered to scaffolds longer than 10 kb, and identical ones dereplicated using CD-HIT v.4.8.1 [68] with settings –c 1.0 –M 100 –n 5. Dereplication was further refined by VIRIDIC v1.0_r3.6 [69] using a 95% intergenomic similarity threshold to define viral species or vOTU (viral operational taxonomic unit) boundaries. The longest or most complete scaffold from each species cluster was kept. To further assess viral genome quality and completeness, CheckV v.1.0.1 [70] was applied.

### Mapping, viral abundance, host prediction, population structure and functional analysis

Host prediction for vOTUs was carried out with iPhop v.1.3.3. with default settings [71] with database Aug_2023_pub_rw. Quality-filtered reads were aligned to vOTUs using Bowtie2 v.2.5.4 [72] with settings –-ignore-quals –-mp 1,1 –-np 1 –-rdg 0,1 –-rfg 0,1 –-score-min L,0,-0.1 –1 [73] to map 90% identical reads according to standard viromics practices [74]. Shrinksam (https://github.com/bcthomas/shrinksam) was used to remove unmapped reads. Average depth of coverage and breadth of coverage (min. 75% for a vOTU to be considered present) were calculated using calc_coverage_v3.rb [75] and calcopo.rb [76] scripts, respectively. Coverage values were normalized to sequencing depth, and relative abundance was calculated as the percentage of each normalized coverage relative to the total. Strain-level diversity assessment was done using inStrain v.1.9 [77] after conversion of .sam files to sorted .bam files using SAMtools v.1.17 [78]. Viral clustering was performed using vConTACT2 v.0.11.3 [79] after gene prediction using Prodigal v.2.6.3 [80], and a taxonomic reference database from INPHARED resource [81], release from January 2025, and further refined by graphanalyzer v.1.6.0 [82] (operating under default parameters). A viral proteomic tree was constructed using VipTree v.4.0 webserver [83] using “Any host” as host category and dsDNA as nucleic acid type for reference viruses. For the ViPTree analysis, a subsample of 120 dereplicated vOTUs classified by CheckV as “medium-quality”, “high-quality”, or “complete” was used.

### Viral gene analysis

Next, DRAM-v.py v.1.5.0 [84] in annotate and DRAM.py in distill mode were employed with the default reference databases – including Pfam, VOGDB, KEGG – to annotate predicted genes with functional assignments. Following the concept by Hurwitz, et al. [85], AVGs were classified into Class I, directly involved in core host metabolism (e.g., carbohydrate degradation, nucleotide biosynthesis, energy pathways, and amino acid metabolism), and Class II, which may indirectly affect host metabolism through functions such as transport, motility (flagellin), peptidases, and vitamin/cofactor biosynthesis. Transport-related AVGs were grouped into subcategories to enhance visualization: amino acid transport (including taurine and peptides), sugar/carbohydrate transport (xylose, glucose, ribose, mannose), inorganic ion transport (sulfate, sulfonate, phosphate, nitrate), and LPS/toxin transport (lipopolysaccharides, toxins, hemolysins).

### Binning and defense systems

The bacterial genomes assembled metagenomes (MAGs) were recovered using MaxBin2 v.2.2.4 [86], and the quality was verified with CheckM v.1.0.13 [87] (Table S1). The bacterial MAGs with >70% completeness and <10% contamination were manually curated using Anvi’o v.8 [88] and dereplicated using dRep v.3.5.0 [89]. The taxonomic assignment was done with GTDB-Tk version 2.1 [90] against database 220 (R220). MAGs were functionally annotated with DRAM v.1.5.0 [84]. Spacer sequences were obtained from CRISPR arrays, which form part of the adaptive immunity of prokaryotes against mobile genetic elements [17]. Only arrays with confidence level 4 were considered for spacer extraction using CRISPRcasFinder [91] in metagenome-assembled genomes and matched to vOTUs (>10 kb) and plasmids (>5 kb, identified by GeNomad) in the sample using BLASTn-short v2.14.0 with a subsequent filtering step of 80% similarity. DefenseFinder v2.0.0 [92] was used to predict the number and type of defense systems in the MAGs. GraphPad Prism v.10.5.0 was used for processing the data and for statistical analysis (D’Agostino & Pearson test was performed to analyse normality, followed by a Welch’s t-test).

### Characterization of carbonic anhydrase

BLASTn and BLASTp analysis [93] of viral-encoded genes functionally annotated as carbonic anhydrase (CA, <u>EC:4.2.1.1</u>), determined from the tree clades, were performed against the Ocean Gene Atlas v.2.0 [94, 95] metagenomic dataset Tara Oceans Microbiome Reference Gene Catalog v1 (OM-RGC_v1) using default settings to check for the CAs’ distributions in the marine environment as well as for microorganismal homologs. The CAs from the vOTUs were further subjected to domain analysis using InterProScan v.5.76-107.0 [96]. To identify additional viral sequences encoding a CA, the InterPro entry of the CA domain (IPR036874) was queried against all proteins in the UniProt database [97]. The CA sequences were aligned using CLUSTAL W [98], visualized using ESPript 3.0 [99], and subjected to phylogenetic analysis using NGPhylogeny.fr [100] with default parameters. An unrooted tree was prepared from the output of the tool NGPhylogeny.fr using Interactive Tree Of Life (iTOL) v7 [101]. Structural characterisation of the CAs was performed using AlphaFold 3 [102], and the ‘model_0’ structures were visualized and aligned using PyMOL v2.4.2. The structures of MAG and related viral CAs were aligned with the vOTU CA_04_176427 (Table S9), while the structures of MAG and *Metania* sp. CAs were aligned with the MAG CA, CA_05_bins_070_1-contigs_c_000000098230_19 (Table S10). Accession IDs, amino acid sequences, confidence of structural prediction, RMSD scores and colour codes are listed in Tables S9 and S10.

### Diversity analysis and statistics

The Shannon-Wiener Index (alpha-diversity) was analysed using the estimate_richness function, while beta-diversity and NMDS analysis based on Bray-Curtis dissimilarity were conducted using the ordinate function, all within the *phyloseq* package v.1.48.0 [103] in the R programming environment v.4.4.0 [104] running on RStudio v.2025.05.0+496. NMDS and beta-diversity plots were generated with the *ggplot2* package v.3.5.2. [105]. Differences in beta diversity between water and sponge tissue, as visualized in NMDS plots, were evaluated using Permutational Multivariate Analysis of Variance (PERMANOVA, *n* = 999 permutations) and betadispersion analysis. These analyses were performed using the adonis2 and betadisper functions, respectively, from the *vegan* package v.2.6-10 [106] in R. A Venn diagram was constructed to check overlaps between sponge and water-associated vOTUs using the UGent webtool (https://bioinformatics.psb.ugent.be/cgi-bin/liste/Venn/calculate_venn.htpl). A virus was considered present in sponge or water, if it was present based on read mapping in one of the four samples.

## Declarations

### Ethics approval and consent to participate

Not applicable.

## Consent for publication

Consent for publication of the photo in Fig.1b is available.

## Availability of data and material

Viral OTUs (full set and 120 vOTU subset) were deposited at figshare [107]. Sequence raw data and MAGs were submitted to NCBI GenBank (https://www.ncbi.nlm.nih.gov/) and can be found under the Project code PRJNA1322391.

## Author contributions

CPPA: sampling, DNA extractions, bioinformatic analysis, editing; RD: bioinformatic and statistical analysis, visualization, writing & editing; OHP: bioinformatic analysis, visualization, editing; GJPJR: bioinformatic analysis, editing; RHK: sampling, funding, supervision, editing; JR: bioinformatic analysis, visualization, supervision, first draft, editing

## Acknowledgements

We acknowledge use of the HPC cluster DRACO, with tools and database kindly provided by the VEO group of Bas Dutilh at the FSU Jena.

