## Supplementary Material for "Viral communities in *Metania* sp. sponge microbiomes with possible effects on CO_2_ fixation"

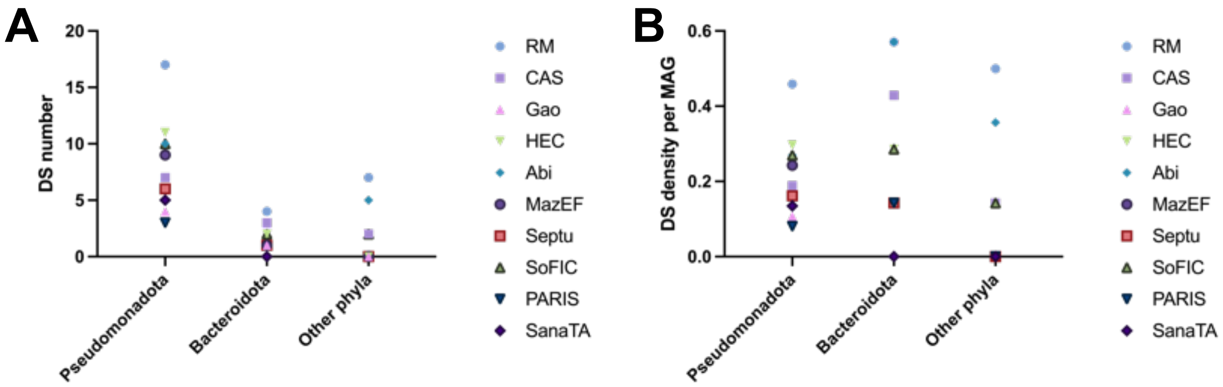


**Figure S1:** Comparison of major DS number among Pseudomonadota, Bacteroidota, and other phyla. Figure B: Comparison of major DS density (per MAG) among Pseudomonadota, Bacteroidota, and other phyla.

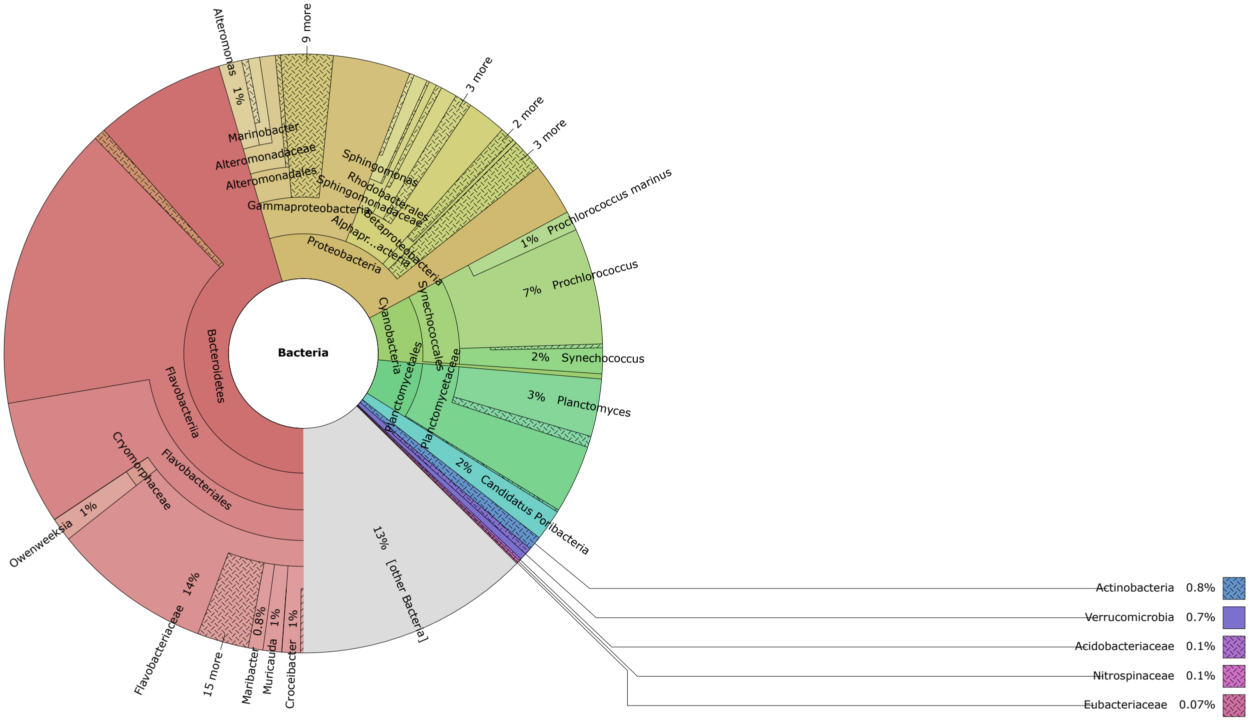


**Figure S2:** Krona plot showing taxonomic distribution of protein homolog of carbonic anhydrase from a vOTU (CA of CA_03_1245020||full16)

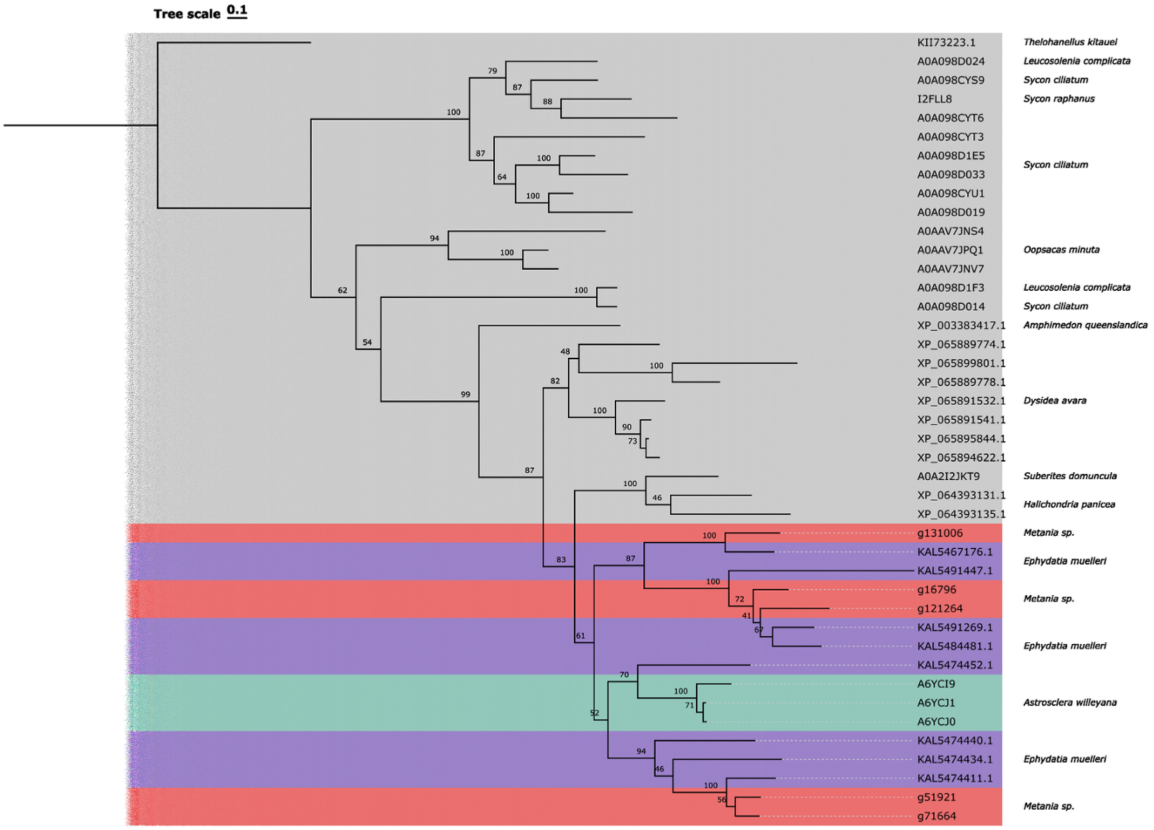


**Figure S3:** Phylogenetic reconstruction of carbonic anhydrase (CA) genes in *Metania* sp. The *Metania* sequences were searched against the NCBI nr database (E-value<1e$^{-30}$). Significant hits (identified by their NCBI accessions), together with the *Metania* sequences, were aligned using Clustal-omega v1.2.2 [1] using the CA signature domain (PF00194) as a guide. A CA representative from *Thelohanellus kitauei* (Cnidaria) was used as the outgroup. Maximum-likelihood phylogenetic tree estimation was performed using IQTREE v3.0.1 [2] with the best-fit model Q.PFAM+R4 and 1,000 ultrafast bootstrap replicates. The consensus tree was rendered using the tvBOT web application[3].
